# Engineering of Primary Human B cells with CRISPR/Cas9 Targeted Nuclease

**DOI:** 10.1101/290973

**Authors:** Matthew J. Johnson, Kanut Laoharawee, Walker S. Lahr, Beau R. Webber, Branden S. Moriarity

**Author notes:** Corresponding Author: Branden S. Moriarity, Cancer and Cardiovascular Research Building, 2231 6^th^ St. SE, University of Minnesota, Minneapolis, MN 55455, USA.

## Abstract

B cells offer unique opportunities for gene therapy because of their ability to secrete large amounts of protein in the form of antibody and persist for the life of the organism as plasma cells. Here, we report optimized CRISPR/Cas9 based genome engineering of primary human B cells. Our procedure involves enrichment of CD19^+^ B cells from PBMCs followed by activation, expansion, and electroporation of CRISPR/Cas9 reagents. We are able expand total B cells in culture 10-fold and outgrow the IgD+IgM+CD27- naïve subset from 35% to over 80% of the culture. B cells are receptive to nucleic acid delivery via electroporation 3 days after stimulation, peaking at Day 7 post stimulation. We tested chemically modified sgRNAs and Alt-R gRNA targeting CD19 with Cas9 mRNA or Cas9 protein. Using this system, we achieved genetic and protein knockout of CD19 at rates over 70%. Finally, we tested sgRNAs targeting the AAVS1 safe harbor site using Cas9 protein in combination with AAV6 to deliver donor template encoding a splice acceptor-EGFP cassette, which yielded site-specific integration frequencies up to 25%. The development of methods for genetically engineered B cells opens the door to a myriad of applications in basic research, antibody production, and cellular therapeutics.

## Introduction

B cells and their downstream effectors, plasma blasts and plasma cells, are central to the humoral arm of the adaptive immune system and are the only cell lineage that secretes antibodies^1, 2^. These antigen-specific antibodies help to protect the host from infection via neutralization and opsonization of pathogens and toxins^3, 4^. B cell are also considered professional antigen presenting cells (APCs) with the ability to present exogenous antigen to naïve T cells on MHC class II^5^. Additionally, B cells play a critical role in the development and maintenance of immunological memory through the generation of memory B cells capable of rapidly reinitiating an antigen-specific immune response upon reencountering their cognate antigen, as well as long-lived plasma cells, which passively maintain low levels of antigen-specific antibodies in the plasma and mucosal surfaces^6^.

Several unique features make B cells and plasma cells an attractive target for genome engineering. B cells are easily isolated in large number from the peripheral blood and can be activated, grown, and expanded in culture^7–9^. Plasma cells specifically upregulate multiple pathways to increase their ability to produce massive amounts of protein and receive pro survival signals from bone marrow stromal cells to extend their longevity^10^. Thus, a system where B cells are isolated from the peripheral blood, engineered to express a specific gene, matured to plasma cells, and then reintroduced back to the host would be an excellent gene therapy platform for long term, high titer expression of protein in the serum.

Previous work on genome engineering B cells has focused on editing primary human peripheral B cells, or editing hematopoietic stem cells and then maturing them to B cells *in vitro*, to express a specific monoclonal antibody (mAb) using lentiviral vectors or transposons^11, 12^. However, the non-site specific integration of lentiviral vectors and transposons and their predilection for integrating into transcriptionally active regions carries the inherent risk of harmful mutations which greatly limits any future clinical applications of such approaches^13^.

The site-specific nature of the CRISPR/Cas9 nuclease system makes it ideal for precision engineering of B cells and recent advancements in CRISPR/Cas9 technologies have drastically improved editing efficiency in primary human cells^14^. The use of high quality Cas9 mRNA or protein and chemically modified guide RNAs (gRNAs) were demonstrated to increase the editing efficiency to over 75% in primary cell types^14^. Additional work has demonstrated that Adeno Associated Virus Serotype 6 (AAV6) functions as a highly efficient DNA template donor for homologous recombination (HR) in both hematopoietic stem cells and T cells^15, 16^. Therefore, we have developed methods for using the CRISPR/Cas9 system with chemically modified sgRNA to knockout endogenous genes and combined this with AAV6 to engineer primary human B cells.

Here we report the site-specific engineering of primary human B cells using the CRISPR/Cas9 system. The use of chemically modified sgRNA is compared with the two-component AltR system^17, 18^ and the use of purified Cas9 protein is compared with chemical modified Cas9 mRNA. Finally, we demonstrate the use of Cas9 protein, AAVS1 sgRNA, and AAV6 as a DNA donor for HR to introduce a DNA sequence encoding splice acceptor-EGFP into the AAVS1 locus of B cells.

## Martials and Methods

### Peripheral Blood Mononuclear Cells (PBMCs) isolation

Human PBMCs from de-identified, normal, healthy donors were obtained by automated leukapheresis (Memorial Blood Centers) and further depleted of red blood cells by lysis with ACK buffer (Thermos Fisher Scientific) for 3 minutes at room temperature. PBMCs were then cultured in RPMI 1640 (Thermos Fisher Scientific) supplemented with 2mM L-gluatmine (Invitrogen), 1% streptomycin and penicillin (Invitrogen), and 10% Fetal Bovine Serum (Glibco) at a density of 1×10^6^ cells/ml. Signed inform consent was obtained from all donors and the study was approved by the University of Minnesota Institutional Review Board (IRB study number 1602E84302). All methods were performed in accordance with relevant the guidelines and regulations.

### Isolation and expansion of B cells

B cells were isolated from PBMCs by immunomagnetic negative selection using EasySep Human B Cell Isolation Kit (Stemcell Technologies) in accordance with the manufacturer’s instructions. B cells were cultured in StemMACS HCS Expansion Media XF (Miltenyi Biotec) supplemented with streptomycin and penicillin (Invitrogen), 5% Human AB Serum (Valley Biomedical), and 125 IU/mL of IL4 (Miltenyi Biotec) at a density of 5×10^5^ cells/ml. B cells were expanded by crosslinking CD40 using Human CD40-Ligand Multimer Kit (Miltenyi Biotec) at a concentration of 8 U/ml in accordance with the manufacturer’s instructions. Media, IL4, and multimeric CD40L were refreshed every 3-4 days throughout the duration of all experiments.

### Electroporations

For electroporations, 3×10^5^ B cells were added to a combination of 1 μg of sgRNA, 1.5 μg of Cas9 mRNA (TriLink), 1 μg of EGFP mRNA (TriLink), or 1 μg of Cas9 protein (Integrated DNA Technologies) and brought up in 10 μl of T Buffer (Thermos Fisher Scientific) for electroporation in the Neon Transfection System (Thermos Fisher Scientific). Cells were then loaded in 10 μl tips and electroporated in accordance with the manufacturer’s instructions using settings of 1400 volts, 10ms width, and 3 pulses unless indicated otherwise.

### Editing with CRISPR/Cas9

gRNAs targeting CD19 and AAVS1 genes were designed using the Crispr MIT website (http://crispr.mit.edu/) along with Cas-OFFinder which together determined the most efficient and specific guides. crRNAs were ordered from Integrated DNA Technologies for testing of the Alt-R system. Tracr and crRNA were combined according to the manufacturers protocol, these cRNA:RNP complexed guides were then mixed with Cas9 protein before being electroporated into primary human B cells^17, 18^. For experiments using chemically modified sgRNAs, guides were ordered from Synthego Biosciences with 2’-O-methyl and 3’ phosphorothioate internucleotide modified linkages at the 3’ and 5’ ends of sgRNAs. sgRNAs were complexed with Cas9 protein for 20 minutes similar to the two component Alt-R system. Guide RNA sequences for CD19 and AAVS1 were CTGTGCTGCAGTGCCTCAA and GTCACCAATCCTGTCCCTAG, respectively. Analysis of Cas9 activity was performed by amplifying a region of genomic DNA around the site of interest. The primer pair AAATTCAGGAAAGGGTTGGAAG and GCGGACCTCTTCTGTCCATG was used to amplify a region flanking the CD19 cut site and primers GAGATGGCTCCAGGAAATGG and ACCTCTCACTCCTTTCATTTGG were used to amply the AAVS1 region. These PCR amplicons were then submitted for Sanger sequencing and subsequent chromatographs were uploaded to the TIDE website (https://tide-calculator.nki.nl) with their respective non-edited control chromatograms^19^.

For experiments inducing a t(8;12) translocation, gRNAs targeting known c-Myc and IGH Burkitt Lymphoma translocation regions were designed^20^. The gRNA sequence was ATAAAGCAGGAATGTCCGAC used for c-Myc and ATATTCCACCCAGGTAGTGG was used for IGH. The primer pair AACCAGGTAAGCACCGAATCC and CTGGCTCACACAGGCGATATGC was used to amplify a region flanking the c-Myc cut site and the primer pair AGCATCTCAATACCCTCCTCTTGG and CAGCCCCAGTTCAGCCTTGTTTAG was used to amply the IGH region and the TIDE website was used to quantify indel formation. The CTGGCTCACACAGGCGATATGC c-Myc reverse primer and AGCATCTCAATACCCTCCTCTTGG IGH forward primer which flank the translocation site were used to confirm the t(8;14) translocation had occurred.

### B cell subset analysis

PBMCs or sorted B cells were washed with PBS and incubated with Fixable Viability Dye eFluor780 (eBioscience) for 10 mins at room temperature. Cells were then wash with a buffer containing 0.5% BSA and stain with fluorescently labeled antibodies to CD10 (Biolegend), CD19 (BD Horizon), CD20 (Biolegend), CD21 (BD OptiBuild), CD27 (Biolegend), CD38 (BD Horizon), CD138 (Biolegend), CXCR5 (BD Horizon), IgD (BD Pharmingen), IgG (Biolegend), and IgM (Biolegend) (supplementary table 1) for 20 mins at room temperature. Samples were then washed a buffer containing 0.5% BSA, fixed with 1% PFA and analyzed a on LSRFortessa flow cytometer (BD Biosciences). Data analysis was performed using Flowjo version 9.9.3 (FlowJo LLC).

### rAAV6 construction and transduction

Splice acceptor-EGFP rAAV virus with 500 base pair homology arms targeting the AAVS1 was produced using custom ordered gBlock Gene Fragment (Integrated DNA Technologies), the AAV backbone, and Gibson assembly. After the vector was constructed it was sent to Vigene Biosciences for AAV6 packaging and concentration. Primary human B cell cultures were transfected with AAVS1 targeting sgRNA pre-complexed to Cas9 protein immediately before being transduced with AAV6 at the indicated MOIs.

### Statistical Analysis

Data analyses were performed using FlowJo version 9.9.3. Statistical analyses were performed using the Paired T test with GraphPad Prism version 7.0c software (GraphPad Software). Error bars represent standard error in all cases. Measurements were considered statistically different at p < .05.

### Data Availability

All data generated and analyzed this this study are included in this published article and supplementary files.

## Results

### Naïve B cells are activated and expanded in culture with CD40L and IL4

To determine the optimal timing and culture conditions for transfections, sorted CD19 B cells were placed into culture and activated by CD40L crosslinking and IL4 for 18 days^21^. B cell numbers remain roughly constant for 7 days before beginning to rapidly expand (Figure 1A). Similarly, the viability of B cells reduced to below 60% initially on days 1 (57 ± 8% (mean ± SD)) and 3 (56 ± 6%), before rebounding on day 7 (90 ± 2%) and remaining over 85% for the duration of the experiment (Figure 1B).

**Figure 1.**
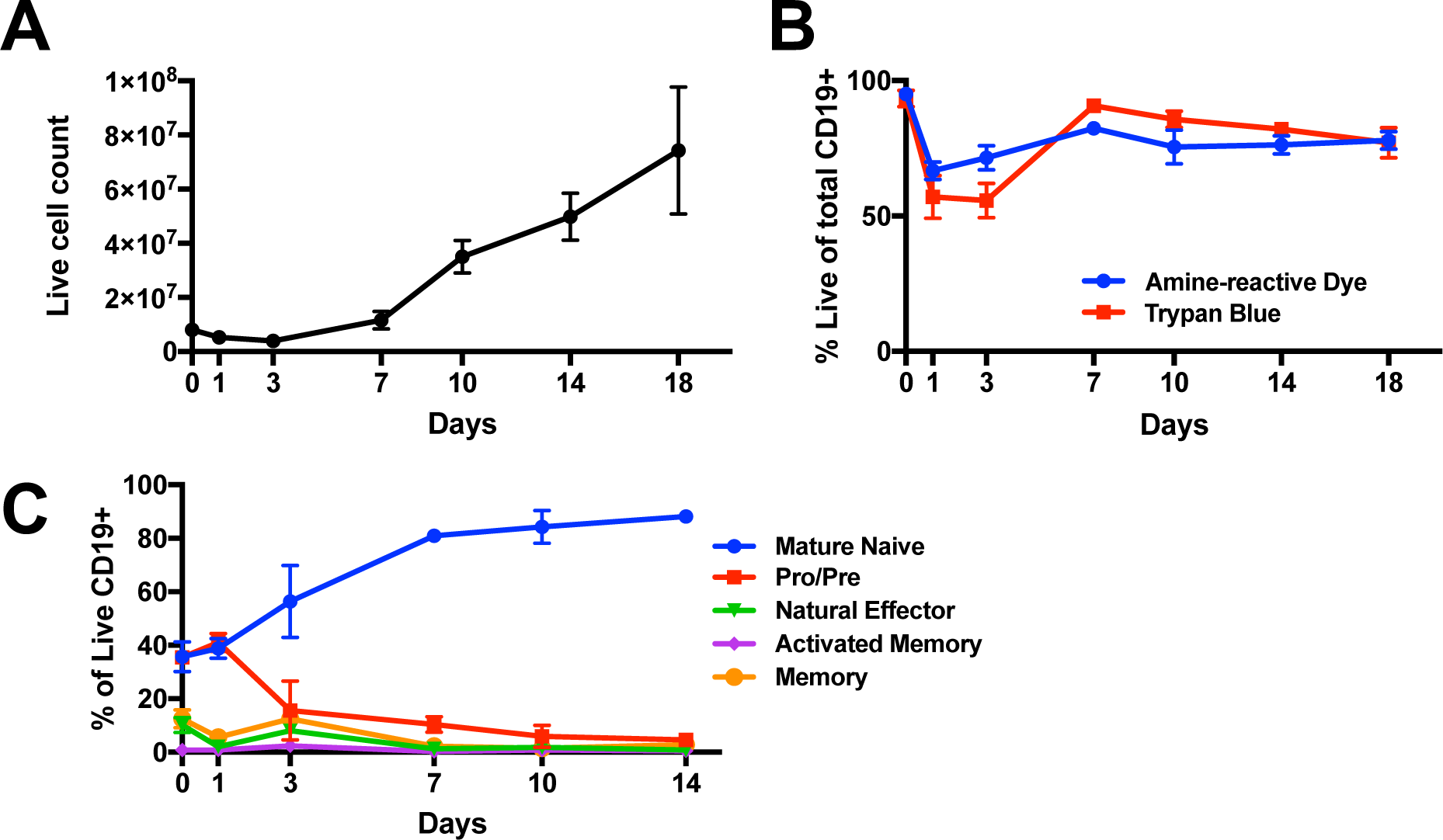
B cells are efficiently expanded with CD40L and IL4. Total cell numbers (A) and viability (B) of isolated B cells in culture and during expansion (n=3). (C) Percentage of Naïve B cells (IgD+ IgM+ CD27-, *blue line*), Pro or Pre B cells (IgD+ IgM- CD27-, *red line*), Natural Effector B cells (IgD+ IgM+ CD27+, *green line*), Memory B cells (IgD- CD27+ CXCR5+, *orange line*), and Activated Memory B cells (IgD- CD27+ CXCR5-, *purple line*) of total live CD19+ B cells in culture over time. (n=3).

Flow cytometry was used to determine the frequency of commonly described subsets of B cells present in the peripheral blood, including IgD+ IgM+ CD27- naïve B cells, IgD+ IgM- CD27- pro and pre B cells, IgD+ IgM+ CD27+ natural effector B cells, and IgD- CD27+ CXCR5+ memory and IgD- CD27+ CXCR5- activated memory B cells (Supplementary figure 1)^22^. Plasmablasts and Plasma cells were undetectable at all time points (data not shown). The naïve B cell population represented 35 ± 6% of the total purified CD19+ B cell culture, but expanded to 81 ± 2% by day 7 and remained over 80% throughout the duration of the experiment (Figure 1C).

### Multiple electroporation conditions allow for efficient transfection of activated B cells

To determine the ideal electroporation condition for CD19+ B cells in culture, we measured the efficiency of EGFP mRNA transfection across 13 conditions of varying voltage, pulse-width, and pulse number (Figure 2A). While all conditions tested achieved over 50% transfection efficiency as measured by GFP expression and over 80% post electroporation cell viability, condition 12 resulted in the optimal combination of EGFP+ frequency, cell viability, and post-electroporation cell expansion (Figure 2B). Therefore, Condition 12, a combination of 1400 volts, 10ms pulse-width, and 3 pulses, was selected for all subsequent experiments.

**Figure 2.**
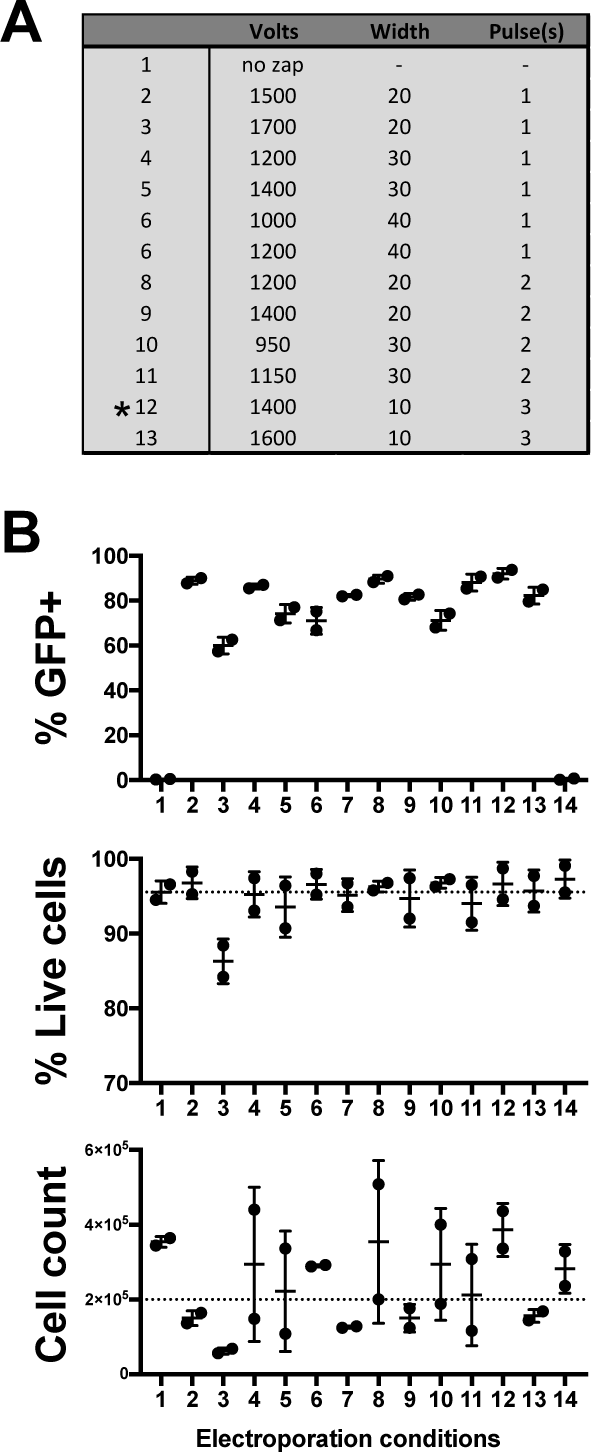
Electroporation conditions affect B cell transfection efficiency, viability, and post transduction expansion. (A) Table of electroporation conditions tested. (B) The percentage of B cells transfected with chemically modified eGFP mRNA *(upper panel)* as well as post transfection cell viability (*middle panel*) and expansion (*lower panel*) (n=2). Dashed lines indicate pre-electroporation viability and cell count.

### B cells become permissive to transfection with mRNA by day 3

To determine the optimal time point for transfection following activation, CD19+ B cells were transfected with mRNA encoding EGFP at multiple time points between day 0 and day 10 post-activation, and the frequency of EGFP+ cells was assayed 48hrs after transfection by flow cytometry (Figure 3). No EGFP expression was detected in any B cell subset following transfection on day 0 and day 1 post-activation, indicating a lack of permissiveness to mRNA transfection at these time points. However, mRNA transfection on days 3, 7, and 10 post-activation resulted in EGFP expression in all subsets. The naïve B cell subset, which at these time points represents the majority of the culture, was transfected at 41 ± 8% on day 3, and over 50% on days 7 (58 ± 9%) and 10 (52 ± 7%). These frequencies are below those observed in Figure 2B, possibly due to reduced cell activation due to the disruption caused by frequent resampling throughout the time-course experiment. All subsequent experiments in this study were performed on Day 7.

**Figure 3.**
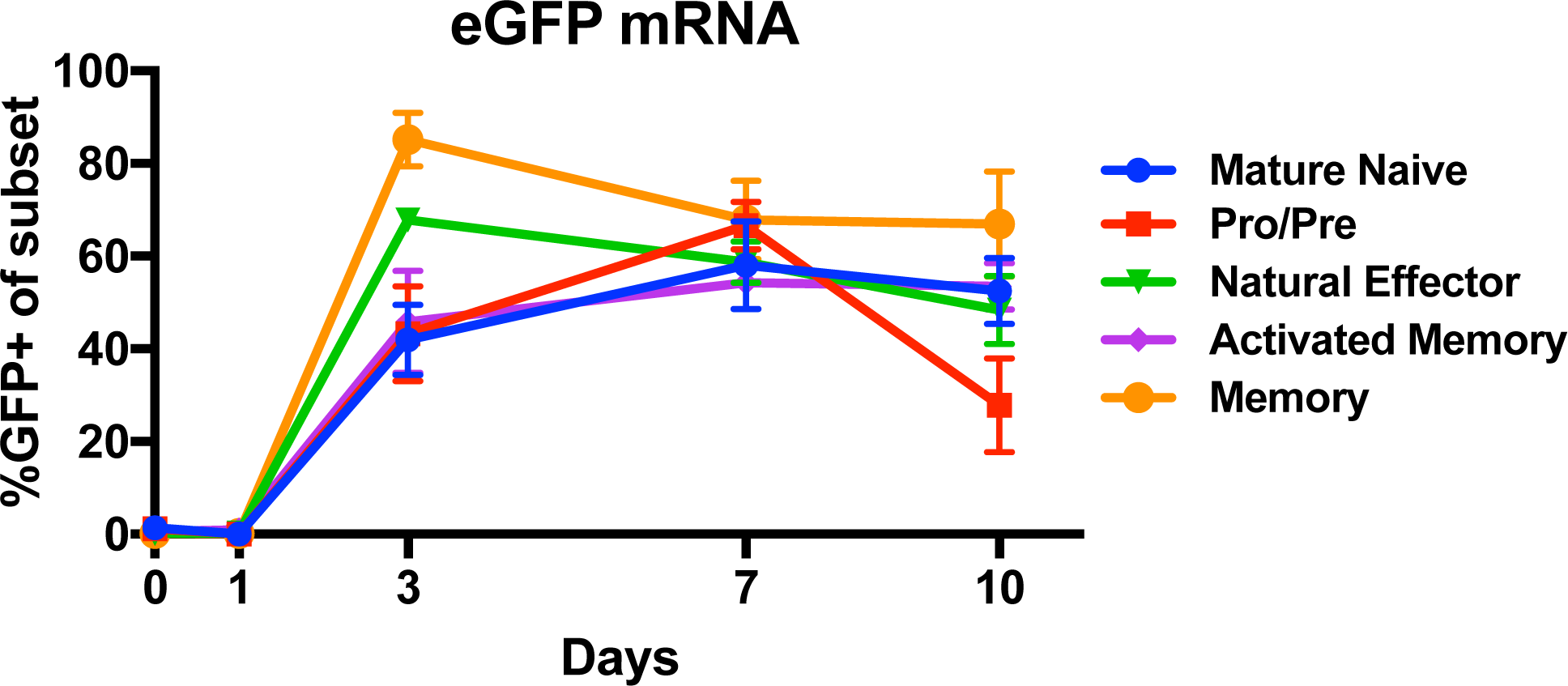
B cells become permissive to electroporation in culture. Frequency of EGFP+ cells 2 days after electroporation with mRNA encoding EGFP (*left panel*) at the indicated time points (n=6).

### Assessment of optimal gRNA and Cas9 format for engineering B cells

Both chemically modified sgRNAs or two component trRNA/crRNA CRISPR/Cas9 systems have been successfully used in primary cells^14, 18, 23^. Therefore, to examine efficacy between two component and sgRNA systems in B cells we targeted CD19, which is expressed on all peripheral blood B cells, with the two component Alt-R system or chemically modified sgRNA in combination with either Cas9 protein or chemically modified Cas9 mRNA (Figure 4A). Chemically modified sgRNA in combination with either Cas9 protein or Cas9 mRNA was able to induce over 70% indel formation (Cas9 protein 74 ± 19% and Cas9 mRNA 72 ± 4%), resulting in >70% loss of cell surface CD19 protein (Cas9 protein 75 ± 11% and Cas9 mRNA 72 ± 13%) in expanded B cells (Figure 4B, *upper panels*). Comparably high editing efficiency was observed with Alt-R gRNA when pre-complexed with Cas9 protein (indel formation 65 ± 9% and CD19 protein knockout 72 ± 11%), however no editing was observed following co-delivery of Alt-R gRNA and Cas9 mRNA.

**Figure 4.**
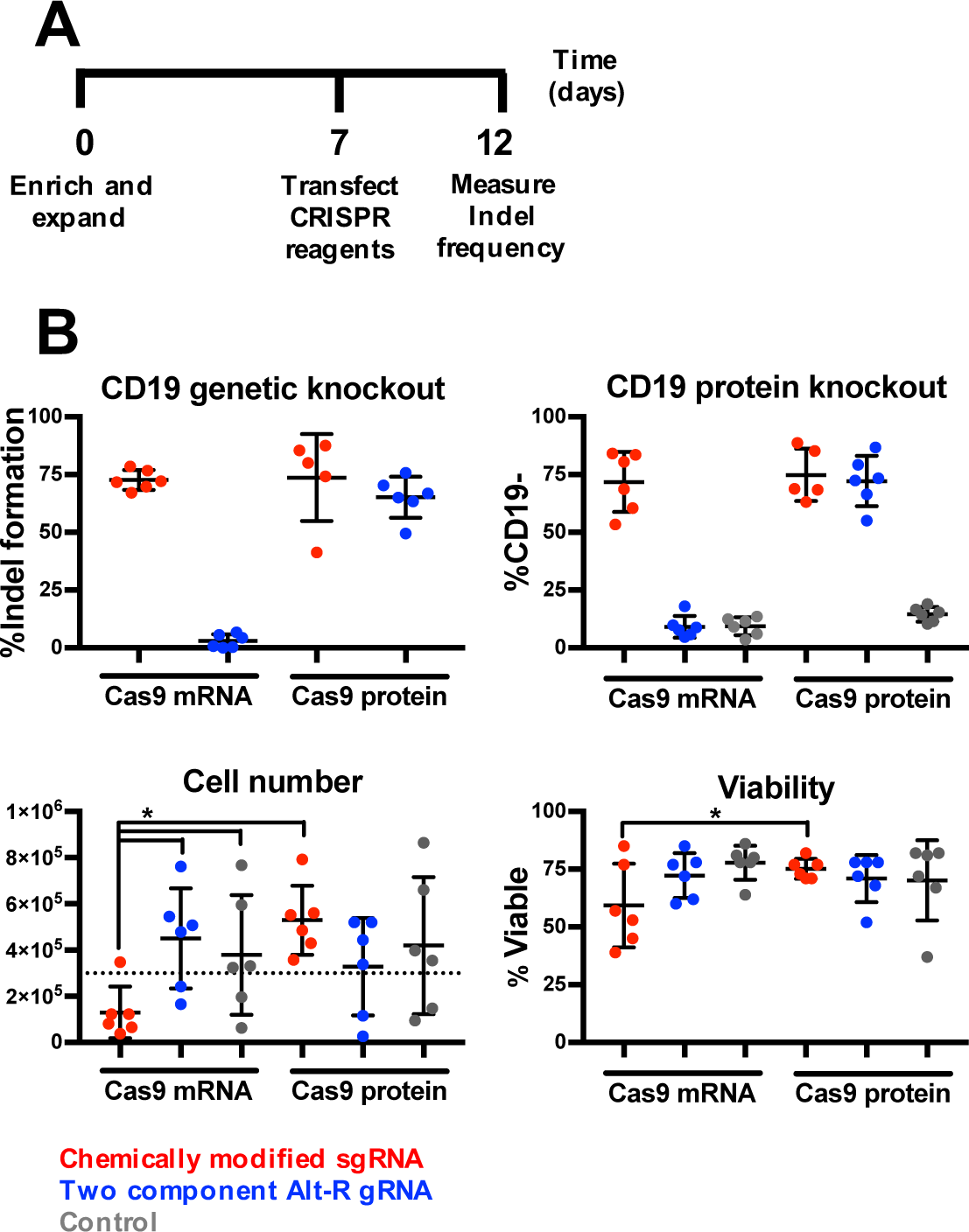
B cell are efficiently edited by CRISPR/Cas9. (A) B cells were enriched and isolated, electroporated with either chemically modified sgRNA or Alt-R gRNA in combination with either Cas9 protein or chemically modified mRNA encoding Cas9 protein, and then assessed for indel formation (n=6). (B) Frequency of indel formation as measured by TIDE analysis (*left panel*) and loss of CD19 expression as measured by flow cytometry (*right panel*), viability (*left panel*) and total cell numbers (*right panel*) (* indicate p <.05) (n=6).

While a high editing frequency was demonstrated in both conditions using chemically modified sgRNA, there were significantly different impacts on the viability of the cells. Cells treated with chemical modified sgRNA and Cas9 mRNA had significantly lower cell numbers (130,000 ± 111,000 cells) and cell viability (59 ± 18%) compared to cultures treated with chemically modified sgRNA and Cas9 protein (529,000 ± 150,000 cells and viability 75 ± 4%) (Figure 4B *lower panels*). In addition, we tested two sgRNAs targeting the BCL2 gene to further validate our findings beyond CD19. When used as part of an RNP these sgRNAs achieved 91 ± 13% and 69 ± 19% indel formation (Supplementary Figure 2). Therefore, the combination of chemically modified sgRNA and Cas9 protein (henceforth referred to as RNP) was chosen for use in all subsequent experiments.

### CRISPR/Cas9 RNPs can be used to induce chromosomal translocations

One potential application of this technology is the induction of chromosomal translocation in a site-specific manner for the study of B cell malignancies. As a proof of concept we modeled the t(8;14) translocation characteristic of most instances of Burkitt Lymphoma (Figure 5A)^24, 25^. In this disease, the gene encoding the transcription factor c-Myc is translocated to the immunoglobulin heavy chain (IGH) locus where enhancers and other elements are thought to strongly up regulate its expression^26–28^. RNPs targeting c-Myc and IGH were used individually and in combination to create DSBs in B cells. Both c-Myc and IGH RNPs created high frequency DSB individual at 78.0% and 56.6% respectively (Figure 5B). As expected, PCR amplification products using c-Myc and IGH locus primer pairs were detectable in all conditions (Figure 5C, *left and center panels*). However, the hallmark t(8;14) translocation, amplified using IGH locus forward primer and the c-Myc reverse primer, was only detected in samples engineered with a combination of RNPs targeting c-Myc and the IGH locus (Figure 5C, *right panel*). The observed band matches the size of the predicted 295bp amplification product.

**Figure 5.**
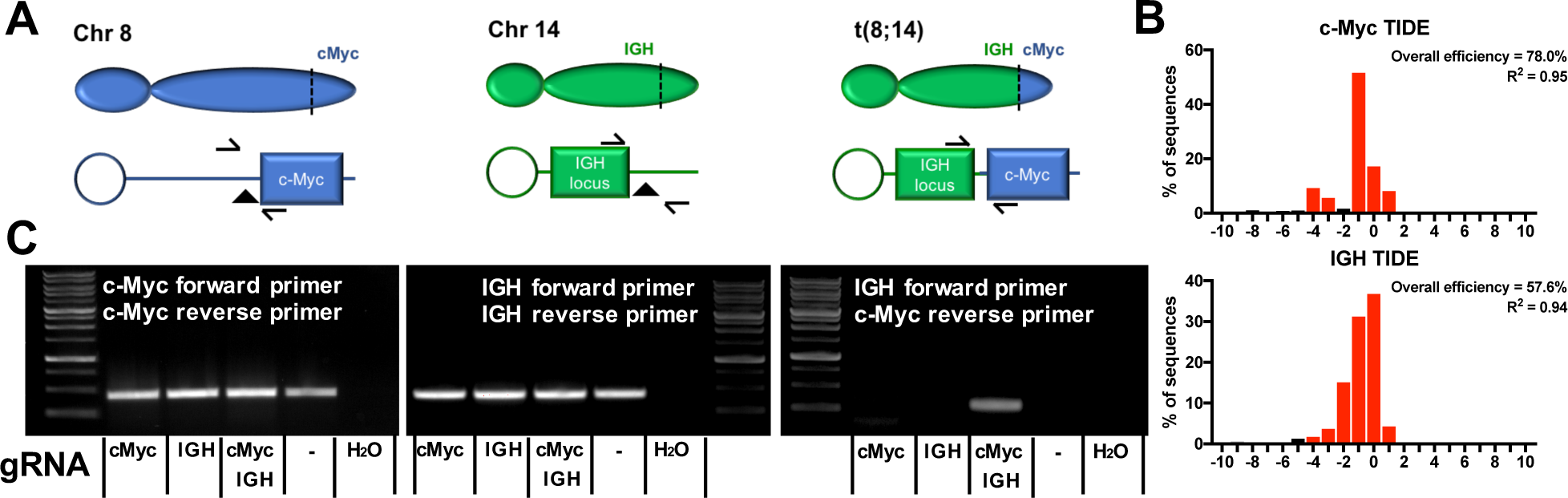
**Simultaneous application of RNPs targeting c-Myc and IGH creates a t(8;14) Burkitt Lymphoma translocation**. (A) Diagram of relative positions of c-Myc on Chromosome 8 (*left diagram*), the IGH locus on Chromosome 14 (*center diagram*) and the t(8;14) translocation (*right diagram*) associated with Burkitt Lymphoma. Arrows indicate primer direction and triangles indicate gRNA target sites. (B) Indel pattern and efficiency of RNPs targeting c-Myc (*top panel*) and IGH (*bottom panel*). (C) PCR amplification using c-Myc (*left panel*), IGH locus (*middle panel*), or a combination of c-Myc/IGH locus primers (*right panel*) of samples engineered with RNPs targeting c-Myc, the IGH locus, or a combination of c-Myc and IGH locus, as well as no gRNA (represented by ‘–‘) and H_2_O controls.

### AAV6 effectively delivers DNA template for HR in B cells

Adeno Associated Virus serotype 6 (AAV6) has been successfully used as a homologous recombination DNA template donor in both CD34+ hematopoietic stem cells and T cells^15, 29^. Given their developmental relationship to B cells as members of the lymphohematopoietic hierarchy, we speculated that AAV6 might also function as a DNA template donor in B cells. To test the efficacy of AAV6 as a DNA template donor, a splice acceptor-based system was designed and constructed that would express EGFP only if integration into the AAVS1 ‘safe harbor’ site was achieved^30, 31^, but not if the vector remained episomal (Figure 6A). Activated primary human CD19+ B cells were electroporated with RNP targeting the AAVS1 locus immediately prior to exposure to AAV6 vector at the indicated MOIs (Figure 6B). Site specific donor DNA integration as evidenced by EGFP expression was observed at all MOIs tested, peaking at 18 ± 4% with an MOI of 250,000. Control samples, transfected with Cas9 protein, but not sgRNA, showed no EGFP expression above background indicating no episomal expression of the EGFP transcript.

**Figure 6.**
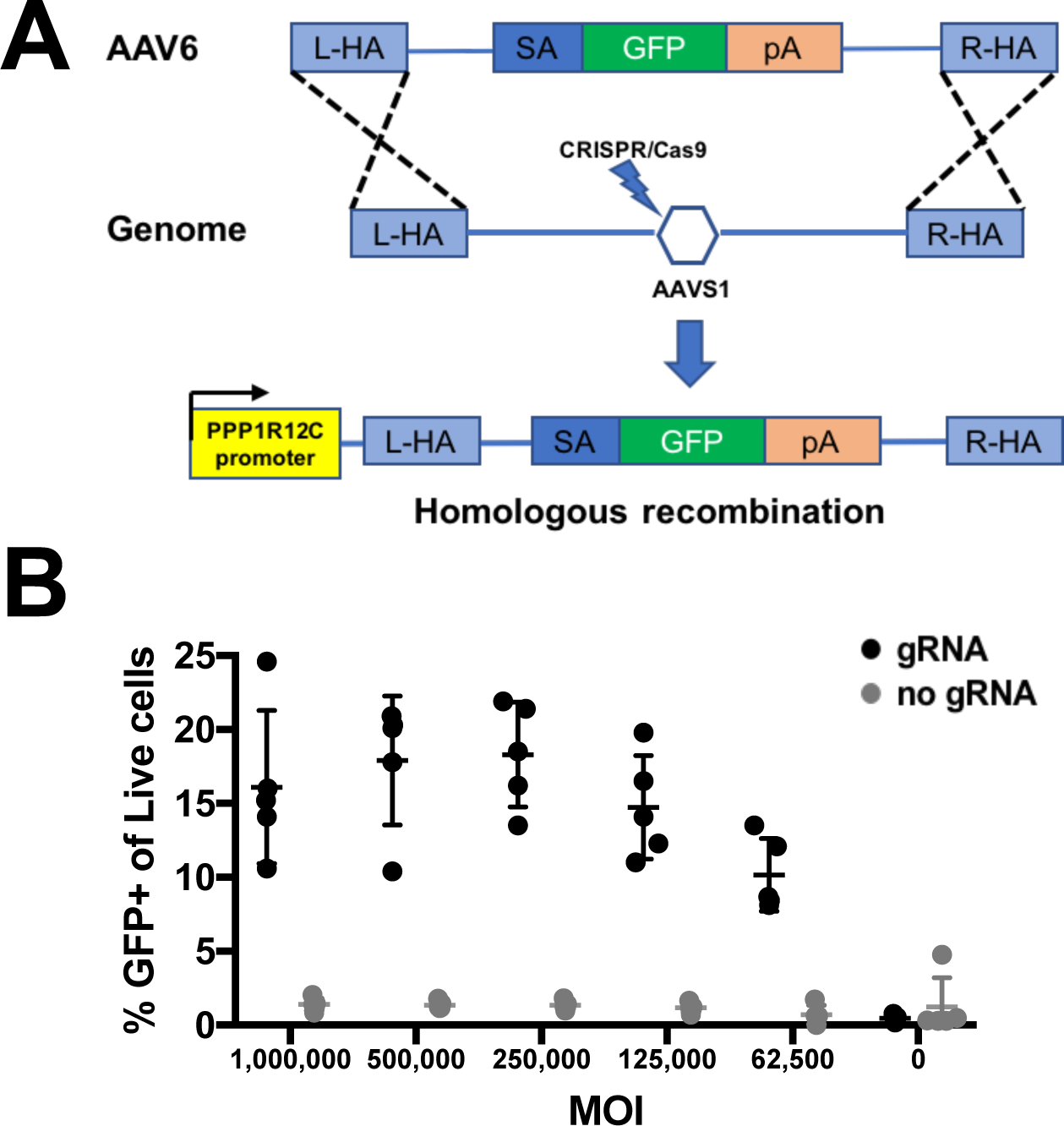
B cells can be engineered using homologous recombination with rAAV6 donor and CRISPR/Cas9. (A) A schematic design of the DNA donor template for a splice acceptor-EGFP system targeting the AAVS1 site by HR used in this study. (B) Percentage of EGFP+ cells represent integration frequency following transfection with rAAV6 containing a splice acceptor-EGFP system at the indicated MOIs (n=3).

## Discussion

B cells are an attractive candidate for use in gene therapy due to their ease of isolation from peripheral blood, rapid growth, and their ability to be converted to long lived plasma cells which produced high levels of protein. The ability to genetically engineer B cells opens a new avenue for using B cells to treat various diseases and to study their basic biology in a human context. Here, we report that RNP, chemically modified sgRNA co-transfected with Cas9 mRNA, and the two-component Alt-R system all efficiently engineer B cells. However, delivery of RNP results in significantly higher post-editing viability and expansion editing. Furthermore, we show serotype AAV6-based donor templates can be successfully used in conjunction with the CRISPR/Cas9 system for targeted integration of a gene of interest at the AAVS1 ‘safe harbor’ site with integration efficiency up to 25%.

While this manuscript was in preparation a group from the University of Washington published a related manuscript on the precision engineering of B cells with CRISPR/Cas9 in *Molecular Therapy*^32^. Many of our overall findings—that B cells can be efficiently edited with CRISPR/Cas9 ribonucleoprotein and that AAV6 is an efficient DNA template donor—are in agreement with their data. In addition, we present data on the subsets of B cells in expanded /activated CD19+ B cell populations, clearly demonstrating that the naïve subset outgrows other subsets in culture and that all subsets are efficiently electroporated after Day 3. We also closely examined alternative methods of CRISPR/Cas9 engineering, demonstrating that the use of Cas9 mRNA, instead of Cas9 protein, has a negative impact on post-engineering B cell health and expansion. This observation may be due to activation of cytosolic RNA sensors such as TLR7, TLR8 or RIG1, which are present in immune cells, including B cells, and have been shown to impact activation, growth regulation, and apoptosis^33, 34^. In additional we demonstrate that CRISPR/Cas9 reagents can be used to model the translocation characteristic of Burkitt Lymphoma. In contrast, their manuscript further describes the *in vitro* maturation of B cells into plasma cells and demonstrates that B-cell activation factor (BAFF) helps to successfully engraft plasma cells into immunodeficient mice. Other, smaller differences in the exact methodologies used, such as differing culture media and B cell isolation kits, appear to have limited impact.

Together, these papers collectively reinforce the viability of using CRISPR/Cas9 engineered B cells as a tool for basic B cell research and as a novel platform for gene therapy. This technology could be adapted for use in research into the basic biology of B cells. Researchers can knockout or overexpress genes for use in both *in vitro* and *in vivo* humanized mouse models. For example, it has been reported that expression of BCL2, BCMA, and MCL1 are essential for the long-term survival of plasma cells *in vivo* in mice^35, 36^. However, their role in the long-term survival of human plasma cells *in vivo* remains difficult to study. CRISPR/Cas9 editing of primary human B cells will make it possible to knockout or over express these gene *in vitro* before maturating cells to plasma cells and adoptively transferring them into a humanized mouse model to assess the gene of interest’s role in *in vivo* plasma cell longevity. Furthermore, we demonstrate the potential for creating chromosomal translocations present in Burkitt Lymphoma. However, an initial characterization of these cell cultures with this translocation showed no obvious transformed properties, i.e. increased proliferation or cytokine independent growth. Moving forward, we will study the process of transformation in greater depth, including determining the impact of additional secondary mutations known to frequently occur in Burkitt Lymphoma^28^.

This technology could also be adapted for the treatment of enzymopathies, a type of genetic order that results in missing or defective enzymes which are usually treated by costly periodic enzyme replacement therapies. Autologous B cells could be isolated, engineered to express the deficient enzyme, and matured into plasma cells before being adoptively transferred back to the original donor where they can express the enzyme. The high protein production potential and longevity of plasma cells could potentially allow for large amounts of the enzyme to collect in the serum for decades after the adoptive transfer. Additional transplants could be applied if serum enzyme levels begin to wane over time.

Treatment using certain monoclonal antibody (mAb) therapies, particularly those used to treat autoimmunity, work by antagonistically binding cytokine or cytokine receptors thereby inhibiting inflammation^37–39^. The biological half-life of these mAbs generally ranges from one to three weeks so biweekly or monthly injections of the mAb are required to keep titers at an efficacious level. Autologous B cells could be engineered to expression therapeutic mAbs, before being matured into plasma cells and adoptively transferred back to the host where they would secrete large amounts mAb into the serum. This could potentially eliminate the need for the repeated injections currently needed in mAb therapies. This could also be further adapted as an alternative method to vaccination for diseases where broadly neutralizing antibodies are protective, but very difficult to elicit using traditional method of vaccination. Further work is required fully realize the potential of the engineered B cell/plasma cell lineage for therapeutic applications.

## Author Contributions

M.J.J., K.L., W.S.L., B.R.W, and B.S.M. designed research; M.J.J., K.L., and W.S.L. performed research; M.J.J., K.L., W.S.L., and B.S.M. analyzed data; and M.J.J. and B.S.M. wrote the paper.

## Acknowledgments

The authors are grateful to Mitchell Kluesner for his careful review of the manuscript.

