## Supplementary Materials for "Engineering of Primary Human B cells with CRISPR/Cas9 Targeted Nuclease"

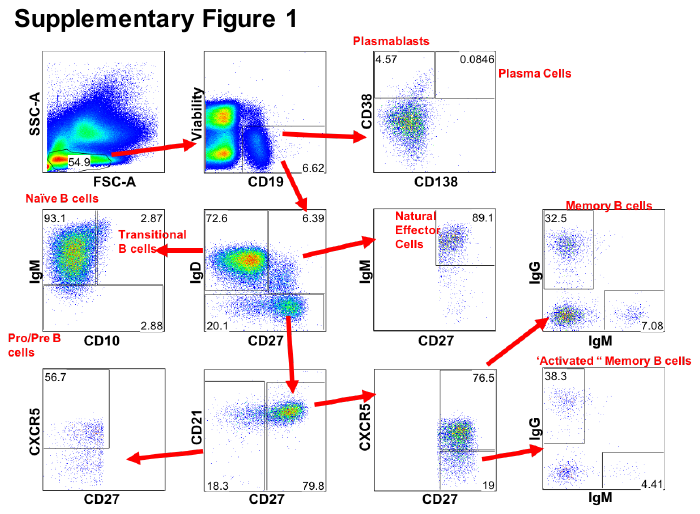

***B cell subsets in the peripheral blood***

Gating scheme used for the identification of B cell subsets by flow cytometry.

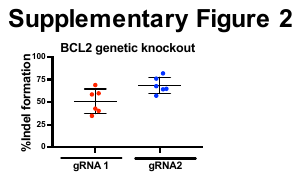

***Knockout of BCL2***

Percent indel formation with two different gRNAs targeting BCL2.

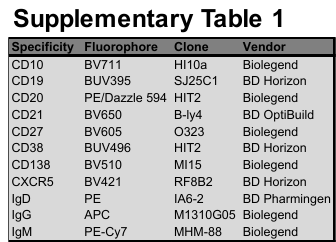

***Flow cytometry antibodies***

List of antibodies used for flow cytometry in this study
